# Marine *Aspergillus terreus* produces a chitinase exhibiting a dual mode of enzymatic action

**DOI:** 10.64898/2026.03.08.710371

**Authors:** Sancharini Das, Debasis Roy, Ramkrishna Sen

**Affiliations:** Department of Bioscience & Biotechnology, Indian Institute of Technology Kharagpur, Kharagpur - 721302, Paschim Medinipur, West Bengal, India; Department of Environmental Science, Savitribai Phule Pune University, Pune, MH, 411007, India; Biochemical Sciences Division, CSIR-National Chemical Laboratory, Pune, MH, 411008, India; Department of Civil Engineering, Indian Institute of Technology Kharagpur, Kharagpur 721302, India

**Author notes:** **Correspondence:** Prof. Ramkrishna Sen, Ph.D., Department of Bioscience & Biotechnology, Indian Institute of Technology Kharagpur, West Bengal 721302, India., Dr. Sancharini Das, Division of Biocehmical Science, CSIR National Chemical Laboratory, Maharashtra, Pune 411008.

**Keywords:** Marine *Aspergillus terreus*, Dual mode of chitinase activity, MALDI ToF/ToF, Peptide finger printing, HRMS, CB Dock 2 molecular docking

## Abstract

Marine *Aspergillus terreus* has been explored as an important chitinase-producing fungal strain for-N-Acetyl-D-Glucosamine (GlcNAc) production from chitin substrates. Here, a purified extracellular 45 kDa chitinase of marine *Aspergillus terreus* (accession number JQ248076) was characterized in terms of substrate specificity. Conventionally, endochitinase cleaves the chitin substrate randomly to produce GlcNAc and its different multimers. So, it requires at least tetramer to characterize the endochitinases; whereas, exochitinases cleaves the chitin substrate from its reducing end and produce either GlcNAc or chitobiose (GlcNAc dimer). In present chitinase characterization, the HPLC followed by HRMS analyses revealed differential product formation from the chitin substrates of varied chain length. With swollen chitin polymer, the enzyme produced GlcNAc as a sole product; whereas with chitohexaose substrate, a mixture of GlcNAc and its oligomers were obtained. Although, mass spectrometry-based proteomics analysis identified the isolated chitinase as an endochitinase 1 precursor (Accession XP_001217186). However, the enzyme kinetic study exhibited higher catalytic efficiency for exochitinase specific dimeric chromogenic substrate in comparison to endochitinase specific tetrameric fluorogenic substrate, which indicated predominantly exochitinase behavior of the enzyme. Further, the in-silico study predicted the differential cleavage pattern of the enzyme, which could be due to different mode of substrate binding and processive mechanism through the tunnel shaped binding cleft of the enzyme. The dual mode of catalytic activity of the present chitinase was further confirmed by a molecular docking study with different lengths of substrates. With the unique dual mode of action, the chitinase of marine *Aspergillus terreus* offers a great promise towards its utility in the production of GlcNAc.

## 1. Introduction

Chitinase (EC 3.2.11.14) is a hydrolytic enzyme that degrades β-1, 4-glycosidic linkage of chitin polysaccharide [1,2], a natural highly abundant biopolymer, second only to cellulose [1,2] and a major component of fungal cell wall and exoskeletal elements of various arthropods, like insects and crustaceans [1–3]. Utilizations of chitinase for N-acetyl-D-glucosamine production from chitin wastes has been widely considered as green method as compared to that of chemical means [1].

Although chitinases are ubiquitous across wide ranges of organisms such as fungi, bacteria, plants, mammals and play diverse physiological role[4,5]. Among all, the fungal chitinases possess unique attributes like production of different chitinase isoforms, unique mode of action including exo-, endo- chitinase activity along with trans-glycosylation activity[6]. Structurally, most of the fungal chitinases belong to the GH 18 family and play many important roles in their cell wall remodeling, morphogenesis, competition, defense and others nutritional purposes[3,7]. Therefore, utility of fungal chitinases on biocontrol, food, agriculture, in pharmaceuticals and in biofuel industries are widely known and well-reviewed [2,8,9]. Intriguingly, chitinases from marine origin exhibit more adeptness in terms of activity against wide physicochemical conditions like salt, pH, temperature etc. [10,11] , thus offer great promise towards several industrial applications.

The most preferred substrate of chitinase is chitin biopolymer, which upon enzymatic degradation produces N-Acetyl -D –glucosamine (GlcNAc) and its oligomers [7,8]. Both GlcNAc and oligomers have industrial and clinical importance [7,8]. On the basis of cleavage pattern chitinases are divided into two major categories – endochitinases and exochitinases. Endochitinases split chitin polymer at internal sites randomly and generate low molecular mass oligomers of GlcNAc [8]. Exochitinases, in contrast are of two types - chitobiosidases (EC 3.2.1.29) and β-(1, 4)-N-acetyl-glucosaminidases (EC3.2.1.30) and excise chitin polymer from its non-reducing end, unlike endochitinases. Chitobiosidases produce dimeric N, N’-diacetylchitobiose whereas, β-(1,4)-N-acetyl-glucosaminidases generate monomeric N-acetyl-D-glucosamine [8]. It was observed that endochitinase activity could be better assessed by using a substrate containing at least a tetramer of GlcNAc. On the other hand, trimer and dimer of GlcNAc are minimal requirement of chitobiosidases and N-acetyl glucosaminidase activity, respectively [12]In recent past, isolation, purification and enzymatic characterization of fungal chitinases have been reviewed in several literatures [7,10,13–15]. A diverse group of fungal species mostly produce cocktail of endo- and exo-chitinases to perform either in cell-wall remodeling or in defense mechanism. Earlier, chitinases isolated from *Aspergillus nidulans* and *Colletotrichum gloeosporioides* exhibited endochitinase activity[14]. Alternatively, β-N-acetylglucosaminidase was found in *Mucor fragilis* and *Trichoderma harzianum* etc.[14]. Intriguingly, air-borne saprophytic *Aspergillus fumigatus* produces a chitinase containing both exo-, endo- and transglycosylation activity [7]. Later, three-dimensional structure demonstrated the structural basis of its unique mode of action[13,14]. Hitherto, many aspects of chitin degradation, structural basis of enzymatic activity have not been investigated fully.

Lately, isolation of marine fungal chitinases, their purification, activities in presence of various metal ions, and antimicrobial potential are discussed elsewhere[10,11]. However, mode of action and structural basis of substrate specificity of the chitinases from marine fungal origin remain obscure.

In this article an extracellular chitinase was purified from a marine *Aspergillus terreus*; which was subjected to undergo enzymatic activity in presence of different substrates with varied length to reveal its mode of action in terms of product formation by natural chitin source. Further biophysical and *in silico* studies including molecular modeling and docking are carried out to understand structural-function relationship in terms of mechanism of action. The current study would provide valuable insights in terms of its utility for GlcNAc production from different chitin sources.

## 2. Materials and Methods

### 2.1. Materials

Swollen chitin was prepared by phosphoric acid and chitin flakes[2]. Glycol chitin was prepared from Glycol chitosan according to Botha et al. (1997) [16]. Hexa-N-acetyl chitohexaose was purchased from Carbosynth, UK. Glycol chitosan, *p*NPGlcNAc, 4-methylumbelliferyl-tri-N-acetyl-β-chitotrioside and Sephadex G-75 were purchased from Sigma Aldrich, USA. All other chemicals were of laboratory or analytical grade and obtained from commercial sources.

### 2.2. Chitinase purification

Marine *A. terreus* was grown in chitinase production medium for three days[2] and the extracellular chitinase was purified from culture medium. The culture filtrate was collected from submerged fungal culture by centrifugation at 4°C and 8000×g and was subjected to salting out by ammonium sulfate precipitation. This was followed by chitin digestion-based affinity chromatography and size exclusion (gel filtration) chromatography. In all the cases the temperature was maintained at 4°C. Initially, culture filtrate (one liter) was subjected to 30% saturation by ammonium sulfate and unwanted biomolecules were precipitated out by centrifuging at 11000×g for 30 minutes. The obtained supernatant was subjected to 70% ammonium sulfate saturation. The desired protein was concentrated in the precipitate upon incubation at 4°C overnight. After centrifuging the precipitate at 11000×g for 30 minutes the protein was resuspended in acetate buffer (50 mM, pH 5.5). The excess salt was removed by dialyzing against same buffer.

Next, to further purify the protein the chitin digestion-based affinity chromatography was set out as reported earlier[12]. In brief, the dialyzed protein (5 mL) was allowed to mix with 1 mL swollen chitin solution (10 mg mL^−1^). The mixture was incubated at 4°C for 30 minutes, letting only chitinase bind chitin matrix, without substantial enzymatic cleavage of chitin polymer at lower temperature[12]. The pellet containing chitin matrix and adsorbed enzyme was separated by centrifugation at 4000×g for 15 minutes at 4°C. The same procedure was repeated with fresh swollen chitin to ensure maximum adsorption of chitinase from the supernatant of earlier step. Pellets from each step were pooled and washed in acetate buffer (50 mM, pH 5.5) supplemented with 1 M NaCl to remove loosely bound unwanted proteins out of the chitin matrix. Protein bound chitin was next re-suspended in 3 mL acetate buffer (50 mM, pH 5.5) and incubated overnight at 37°C to initiate chitin digestion. The digested sample was centrifuged at 4°C and 4000×g for 30 minutes and the chitinase rich supernatant was collected [17]. To further purify, the supernatant was concentrated and loaded onto pre-equilibrated Sephadex G-75 gel filtration resin for the removal of non-specific contaminant. Eluted protein was monitored at 280 nm at a flow rate of 1 mL per minute [12]. Molecular weight of the purified protein was evaluated by running the sample through SDS 12% PAGE [12].

### 2.3. In-plate chitinolytic activity

Chitinolytic activity of the purified protein sample was assessed in chitin-agarose plate. The plates were prepared by mixing 0.5% (w/v) soluble glycol chitin in 1% agarose. Four circular wells were prepared in the chitin-agarose plates where purified protein was loaded into two wells. The other two wells were filled with acetate buffer (50 mM, pH 5.5) and commercially available chitinase from *Trichoderma harzianum* (10mg mL^−1^) as negative and positive control respectively. The sample loaded plate was incubated at 37°C for 3.5 h to allow hydrolysis of glycol chitin. Detection of chitinolytic activity of the sample was carried out by incubating the plates with 0.1% Ranipal solution for 15 minutes. The plates were thoroughly washed by distilled water for at least five times and the zone of clearance was observed under UV transilluminator at 254 nm [17].

### 2.4. Protein identification by tandem mass analysis

To identity of the purified protein, the protein was fractionated by SDS-PAGE (Figure S2.). The protein band was excised from the gel and chopped into small pieces. The gel pieces were destained with solution of ammonium bicarbonate (25 mM) and acetonitrile solution (1:1) at least for three times [18]. The protein inside the gel pieces were reduced by DTT (10 mM) and the cysteins residues were alkylated by iodoacetamide (55 mM) as described earlier[18]. Next gel pieces were subjected to trypsin digestion (10 ng μl^−1^) following published protocol [18].

The extracted tryptic peptides were subjected to Maldi ToF /ToF analyses to obtain peptide mass finger printing. The dried tryptic peptides were reconstituted in trifluoro acetic acid (0.1%) containing acetonitrile solution (50%) and α-Cyano-4-hydroxycinnamic acid matrix was mixed prior to analysis. The matrix mixed peptides were spotted onto MALDI plate and subjected to ionize through laser (4800 MALDI ToF/ToF, Applied Biosystems, Foster City, CA, USA). Significantly intense peaks obtained in the mass spectra were pooled for Collision Induced Dissociation (CID) using air as collision gas. The tandem mass spectra were searched against NCBI non redundant protein database of other fungi (except *Saccharomyces* spp.) using MASCOT software (Matrix Science Ltd., London, UK) to determine protein id with maximum missed cleavage site 2, fixed carbamidomethyl modification and variable methionine oxidation. For MASCOT, the precursor mass tolerance was set to 50 ppm and fragment mass tolerance was 0.6 Da.

### 2.5. Swollen chitin and hexa-N-acetylchitohexaose hydrolysis

To investigate how chitinase enzyme acts on different substrates, the purified enzyme was allowed to enzymatic hydrolysis with swollen chitin and hexa-N-acetyl chitohexaose, individually. A suspension of swollen chitin (1% w/v) allowed to react with purified chitinase (1 mg mL^−1^ in 50 mM acetate buffer pH 5.5) at 45°C for varied time period of 1 h to 24 h. The enzymatic hydrolysis was stopped by immersing the mixture in a 100°C water bath for 5 minutes. The reaction mixture was centrifuged at 5000 g for 10 minutes and the product released in the supernatant was subjected to HPLC and HRMS analyses[9].

Hydrolysis of hexa-N-acetylchitohexaose was carried out by mixing 5 nmol substrate in 10 μl acetate buffer (50 mM, pH 5.5). The reaction mixture was incubated at 45°C for varied time period of 5 minutes to 3 h. The reaction was stopped by boiling the mixture at 100°C [7] and was subjected to product identification by HPLC and HRMS analyses.

### 2.6. HPLC and HRMS analyses

The hydrolysate (20 μl) of hexa-N-acetylchitohexaose and swollen chitin was injected and separated through Zorbax carbohydrate column (250 × 4.6 mm, 5 μm particle size) fitted in high performance liquid chromatography (Agilent Technologies,1100 series, CA, USA). The separation was performed through a linear gradient of acetonitrile and deionized water (95:5) and monitored by diode array detector (Agilent 1100 series DAD). The eluted fractions were pooled and lyophilized prior to its mass analyses by high resolution ESI-MS (HRMS) [19].

The lyophilized powder samples (0.5 mg) were dissolved in acetonitrile solution in water (50%) and analyzed by high resolution mass spectrometer (HRMS) (Xevo G2 QTof, Waters, MA USA) for the determination of mass as well as elementary composition of the hydrolyzed products. The sample was applied to the instrument by infusion method and scanned in both positive and negative ion mode at a mass range of 100 to 1000 Da [2].

### 2.7. Enzyme kinetics

To determine exochitinase activity, 50 μl soluble substrate *p*-Nitrophenyl N-acetyl-β-D-glucosaminide (*p*NPGlcNAc, 5 mM) was allowed to react with 50 μl purified chitinase (0.2 mg mL^−1^) in acetate buffer (50 mM, pH 5.5) with 200 μl of acetate buffer. The reaction mixture (300 μl) was incubated at 45°C for 30 minutes and enzymatic hydrolysis was stopped by 1 mL sodium carbonate (200 mM) solution. The amount of *p*-nitrophenol produced was monitored spectrophotometrically at 405 nm. One unit of exochitinase or N-acetylglucosaminidase activity of the purified chitinase was defined as the amount of enzyme required to produce 1 μmol of *p*-nitrophenol per second. Kinetics parameters of the purified enzyme (*K*_m_, *V*_max_, *K*_cat_ and *K*_cat_/*K*_m_) were determined by performing the set of reactions with varied substrate concentrations (1 to 10 mM) and purified chitinase [20].

To assess endochitinase activity of the purified enzyme an artificial fluorogenic substrate 4-methylumbelliferyl-tri-N-acetyl-β-chitotrioside was used. The reaction mixture (200 μl) was prepared by purified chitinase (10 μl, 0.8 mg mL^−1^) varying substrates concentrations (1 to 10 mM) in acetate buffer (50 mM, pH 5.5). The mixture was incubated at 45°C for 15 minutes. The enzymatic hydrolysis was stopped by the addition of sodium carbonate (100 μl of 200 mM) solution in the reaction mixture. One unit of endochitinase activity was defined as the amount of enzyme required to release 1 μmol of 4-methylumbelliferone (product) per second at 45°C [21]. The release of product was monitored at an excitation of 360 nm and emission of 450 nm through Cary–Eclipse spectro-fluorometer (Varian Inc. Palo Alto, CA, USA).

### 2.8. Statistical analysis

Statistical analysis for all the experiments was performed in triplicates and data were analyzed as mean ± standard deviation. Enzyme kinetics parameters were derived by Graph Pad Prism 5 software. [22]

### 2.9. Prediction of subcellular localization

The presence of signal peptide in the amino acid sequence was predicted by using SignalP 4.1 server (http://www.cbs.dtu.dk/services/SignalP-4.1/) [23]. Subcellular localization predictor (SCLpred) server was used to determine the localization of the chitinase[24]. In both the cases, amino acid sequence submitted to corresponding servers. The amino acid sequence of the protein was fetched from NCBI database with the protein id obtained by MALDI ToF/ToF analyses.

### 2.10. Secondary structure determination

Circular dichroism (CD) spectroscopy and bioinformatics tools were used to determine the secondary structure of the protein.

A far UV circular dichroism (Far UV CD) spectrum of purified enzyme (0.5 mg mL^−1^) was recorded in Jasco J-810 spectropolarimeter (Easton, MD, USA). The sample was loaded in a quartz cell (path length = 0.1 cm) at 25°C and spectral data was collected with step resolution of 0.2 nm, time constant of 1s, sensitivity 10 milli degrees, scan speed of 50 nm/minute and a spectral bandwidth of 2.0 nm. Protein spectrum was corrected by the buffer spectrum and average spectrum of four successive accumulations was collected in a range of 190-240 nm to reduce the noise and random instrumental error. The secondary structures of the purified protein (percentage of α-helix, β-sheet and random coil) were calculated by k2d3 software from CD spectral data[25] .

Different bioinformatic tools were employed to predict the secondary structure of the protein. The obtained amino acid sequence from NCBI database was submitted to PSIPRED v3.3 (https://bioinf.cs.ucl.ac.uk/psipred/) [26]and Predict protein (https://www.predictprotein.org/) [27]servers to determine percentage of α-helix, β-sheet and random coil.

### 2.11. Three-dimensional structures predictions

SWISS–MODEL workspace (http://swissmodel.expasy.org/workspace) was used to build a three-dimensional model of *A. terreus* chitinase [28]. SWISS-MODEL works in fully automated mode with the satisfaction of spatial restraints to build a homology model. Initially, the FASTA format of the target sequences was extracted from the NCBI database and submitted to the aforementioned website to build the model. An automatic template selection was made by the server on the basis of a Blast E-value limit from SWISS-MODEL template library [29] . After that multiple sequence alignment is formed between template and desired protein. The stereo-chemical properties of the generated model are assessed by ANOLEA mean force potential and Gromos 96 force field energy [29]. The stereochemistry and stability of the model were verified by PROCHEK [30] and Verify 3D [31] (http://nihserver.mbi.ucla.edu/Verify_3D) programs.

### 2.12. Molecular docking

*p*-Nitrophenyl N-acetyl-β-D-glucosaminide (PubChem SID 24897896) and 4-methylumbelliferyl-tri-N-acetyl-β-chitotrioside (PubChem SID 24897027) were used as ligands for the molecular docking study of derived *A. terreus* chitinase model. Here, SDF files of both ligands were downloaded from the NCBI-PubChem database and used for docking. CB-DOCK 2 tool (https://cadd.labshare.cn/cb-dock2/php/index.php) was used for docking these ligands with *A. terreus* chitinase receptor. To perform the receptor-ligand docking, the PDB file of the receptor and SDF files of the ligands were uploaded in CB Dock 2 server and blind docking were performed. Further the 3D interaction of the docked complexes was analyzed through PyMOL software [32]

## 3. Results

### 3.1. Chitinase purification

The extracellular chitinase of marine *A. terreus* was purified step by step through ammonium sulfate precipitation, chitin digestion-based affinity chromatography, and size exclusion chromatography. In brief, marine *A. terreus* was grown for three days and the culture filtrate was subjected to salting out by gradually increasing ammonium sulfate concentration. It was observed that the most chitinolytic activity was concentrated at a saturation of 70% ammonium sulfate, which also contained lower contaminants.

The protein fraction from ammonium sulfate precipitation was subjected to further purification through enzyme substrate affinity chromatography, also known as chitin digestion[7]. The chitin-digestion based affinity chromatography was carried out in three steps – binding, wash and elution steps. In binding step, the desired enzyme was allowed to adsorb in swollen chitin matrix at lower temperature. The unbound or loosely bound proteins were removed after thorough wash with buffer in the second step. In the third step, the catalysis was initiated by rising temperature of the protein bound matrix to 37°C, where swollen chitin was degraded, and the enzyme was stripped off the matrix.

Next, gel filtration chromatography was set out in order to further purify the protein eluted from affinity chromatography. The protein was concentrated to a minimal volume and finally loaded onto Sephadex G-75 size exclusion resin and eluted with acetate buffer (50 mM, pH 5.5). The molecular weight of the purified chitinase was around 45 kDa as determined from SDS PAGE analyses (Figure 1).

**Figure 1.**
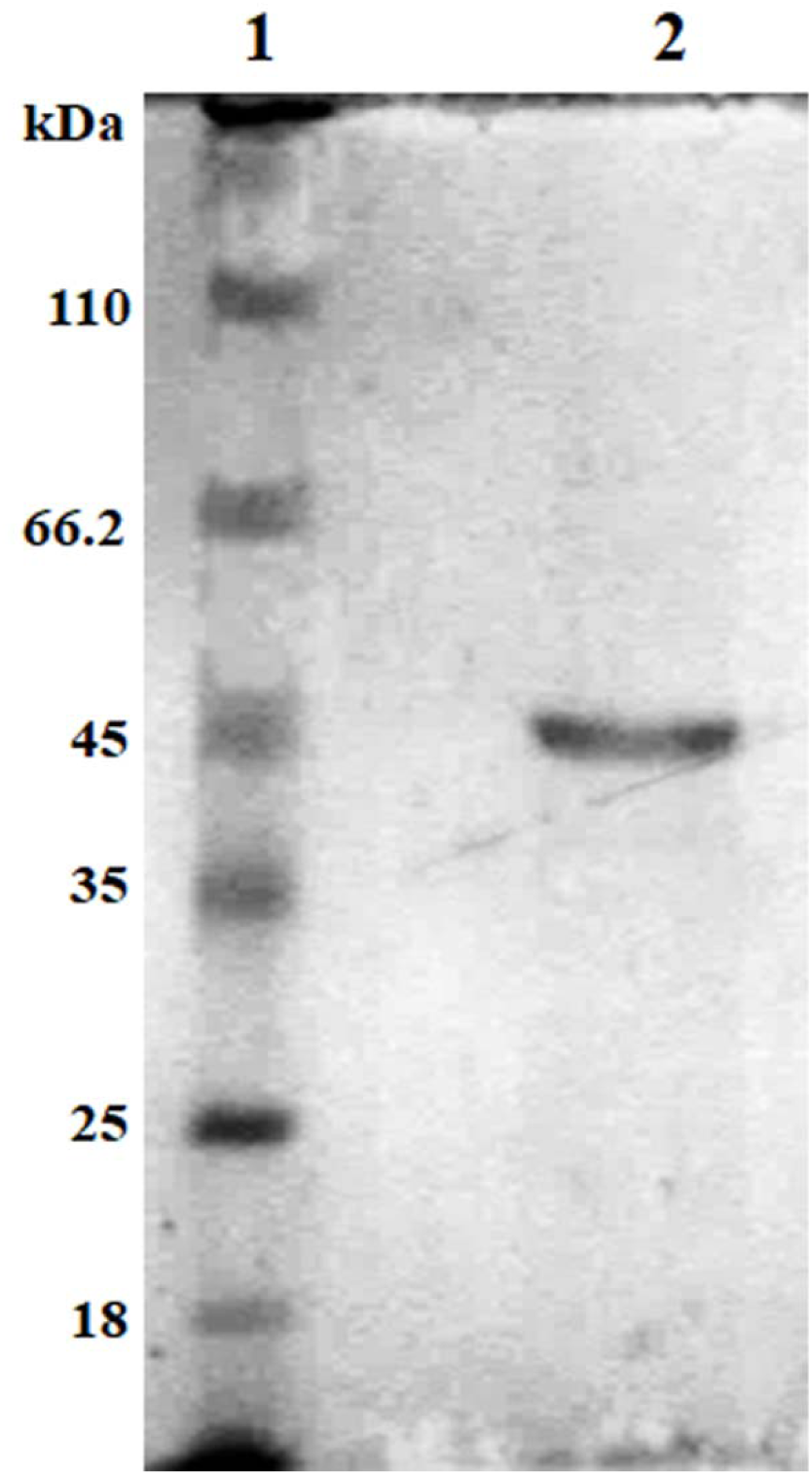
SDS PAGE of purified *A. terreus* chitinase; where, Lane 1: molecular weight marker, Lane 2: Purified *A. terreus* chitinase

The chitinolytic activity was tested at each step of purification and in every eluted fraction. The fold increase in specific activity at different purification steps determined the extent of purity of the protein at each step (Table 1).

**Table 1.** Summary of specific activity after different steps of chitinase purification.

| **Purification step** | **Total volume (mL)** | **Total protein (mg)** | **Total activity (U min<sup>-1</sup>)** | **Specific activity (U mg<sup>-1</sup>)** | **Fold purification** |
| --- | --- | --- | --- | --- | --- |
| Culture filtrate | 1000 | 39 | 12 | 0.31 | – |
| Ammonium sulfate precipitation | 22 | 15.73 | 5.3 | 0.34 | 1.1 |
| Chitin digestion | 5 | 4.83 | 2.8 | 0.58 | 1.87 |
| Gel filtration Chromatography | 3 | 3.22 | 2.35 | 0.73 | 2.35 |

**Table 2.**
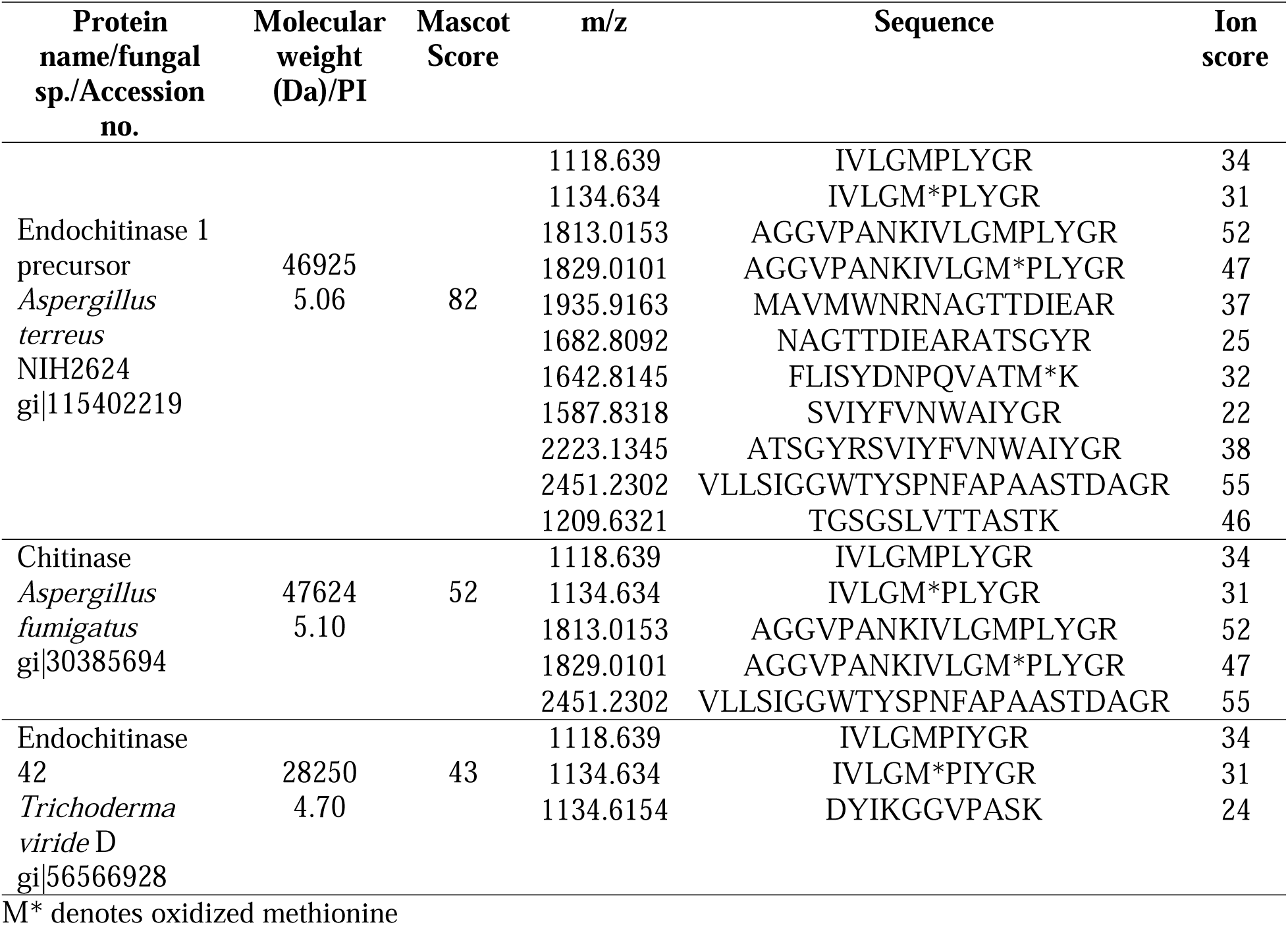
Peptide mass finger prints by Tandem MS/MS analysis of purified chitinase.

### 3.2. Chitinase detection by plate assay

A qualitative detection of the chitinolytic activity of purified chitinase was performed by glycol chitin-based plate assay; where the plate was stained with Ranipal, a diaminostilbene compound containing fluorescent dye and possesses its affinity for binding with the β-glycosidic linkage of the polysaccharide[17] . Therefore, Ranipal binds only with intact chitin molecules, not with the hydrolyzed product. Un-hydrolyzed glycol chitin in the plate-bound with the Ranipal produced intense blue-fluorescent color under UV illumination. In contrast, the chitinase hydrolyzed part around the wells of the plate was observed as a clear zone (Figure S1). The purified chitinase from *A. terreus* exhibited a high level of chitinolytic activity as observed by the zone of clearance on the plate.

### 3.3. Tandem mass analysis of purified chitinase

A combination of SDS PAGE and MALDI ToF/ToF was employed in order to identify the protein profile of the purified chitinase of *A. terreus*. The coomassie stained protein band was excised into small pieces and subjected to peptide-mass fingerprinting analyses. The trypsin-digested peptide masses as well as their fragment ions were determined by MALDI ToF/ToF, which generated nine peptides. The peptide mass spectral data were fed into a MASCOT aided database query, which resulted to the identification of purified enzyme (Table. 2). Identification of the protein was performed by setting A MASCOT score of greater than 60. The protein was identified as endochitinase 1 precursor with accession # XP_001217186 (gi|115402219) (427 amino acids). Molecular mass and PI of the identified protein was 46.9 kDa and 5.06, respectively as determined by MASCOT analyses.

### 3.4. Swollen chitin and hexa-N-acetyl chitohexaose hydrolysis

The proteomics analyses identified the purified chitinase of *A. terreus* as endochitinase 1 precursor. However, there are no biochemical evidence on substrate specificity and cleavage pattern that could categorize the present chitinase to a particular class. Since, appraisal of chitinase enzyme is determined on the basis of type of product formed [10]; therefore, we reasoned that enzymatic characterization of *A. terreus* chitinase in presence of substrates of varied length would presumably evaluate the nature of enzyme in a proper way. In this study a combination of HPLC and HR-MS analyses were performed to identify the products.

In brief, swollen chitin polymer was subjected to hydrolyze and the hydrolysate was subjected to HPLC followed by HR-MS analyses. Both HPLC and mass spectrometric analyses revealed GlcNAc as the sole product of enzymatic hydrolysis reaction of swollen chitin in positive ion mode of the HRMS (Figure 2). This observation led to confirm the exochitinase behavior of purified chitinase of marine *A. terreus*. In contrast, hexa-N-acetylchitohexaose hydrolysis revealed the production of GlcNAc oligomers at different time of incubation. Intriguingly, hydrolyzed products of hexa-N-acetyl chitohexaose were eluted within the retention time 10 to 17 minutes as observed by HPLC choramogram and identified as GlcNAc (Figure 3.a), di-N-acetylchitobiose (Figure 3.b), tri-N-acetylchitotriose (Figure 3.c) and tetra-N-acetylchitotetraose (Figure 3.d) by HR-MS analyses. Presence of Tri-N-acetylchitotriose during enzymatic hydrolysis of hexa-N-acetylchitohexaose indicated the endochitinase behavior of purified enzyme. Thus, the purified chitinase of *A. terreus* exhibited dual mode of action in presence of different substrates of varied size and length.

**Figure 2.**
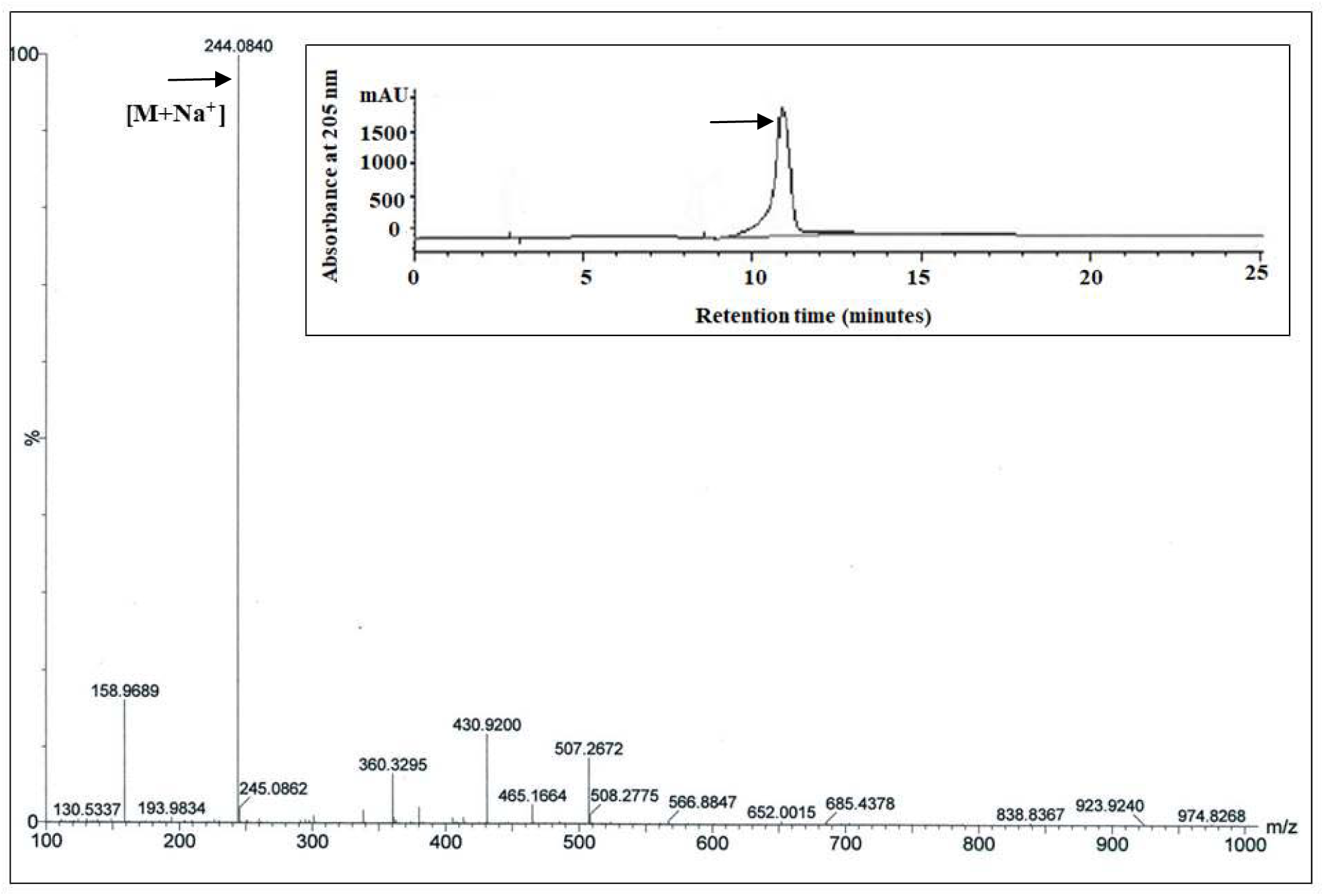
HRMS of the HPLC purified fraction of swollen chitin hydrolysed product. The data was recorded in positive ion mode of the instrument. Sodium ion (Na^+^) adducted GlcNAc was detected as predominant product (HPLC chromatogram is represented in the inset)

**Figure 3.**
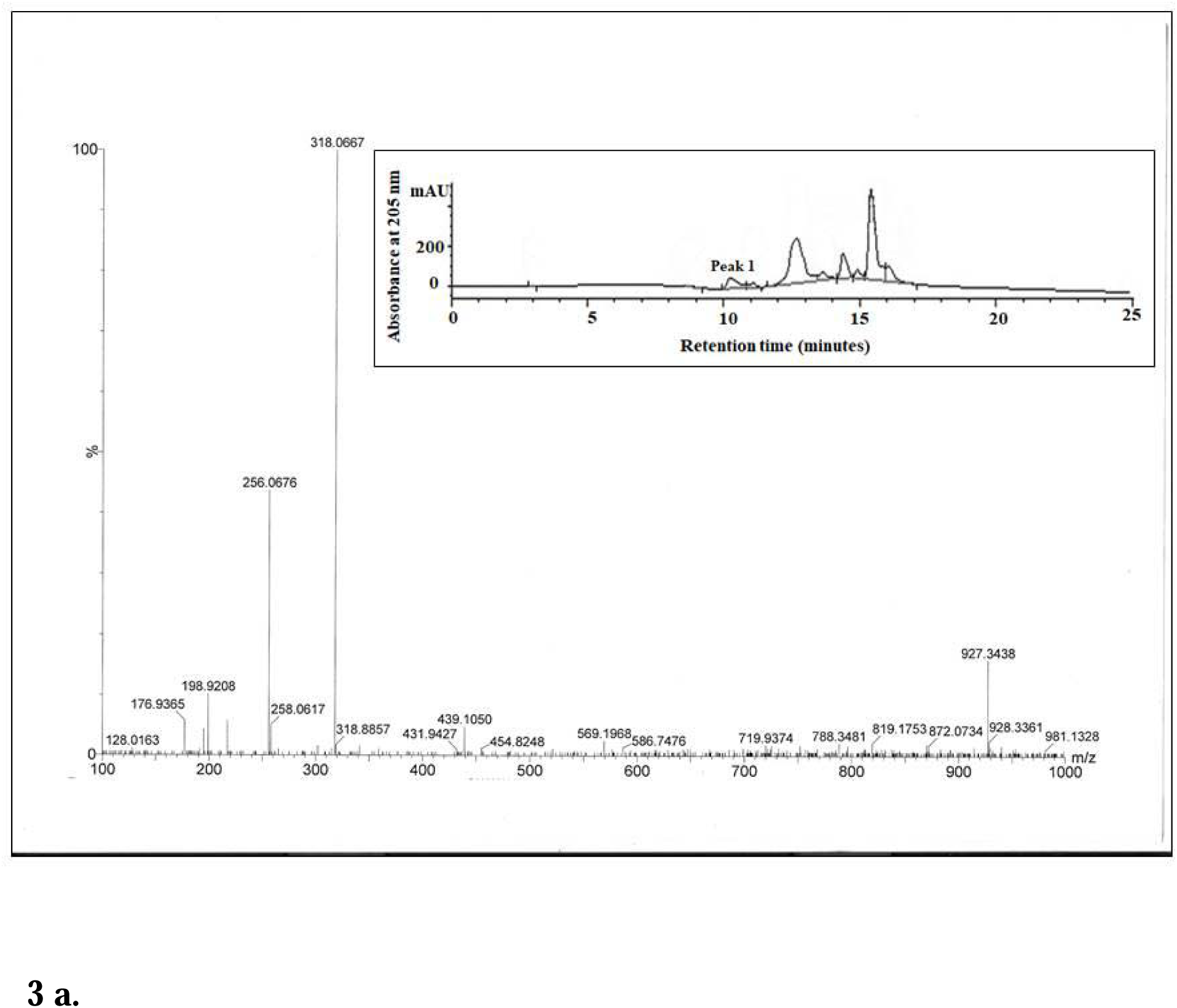
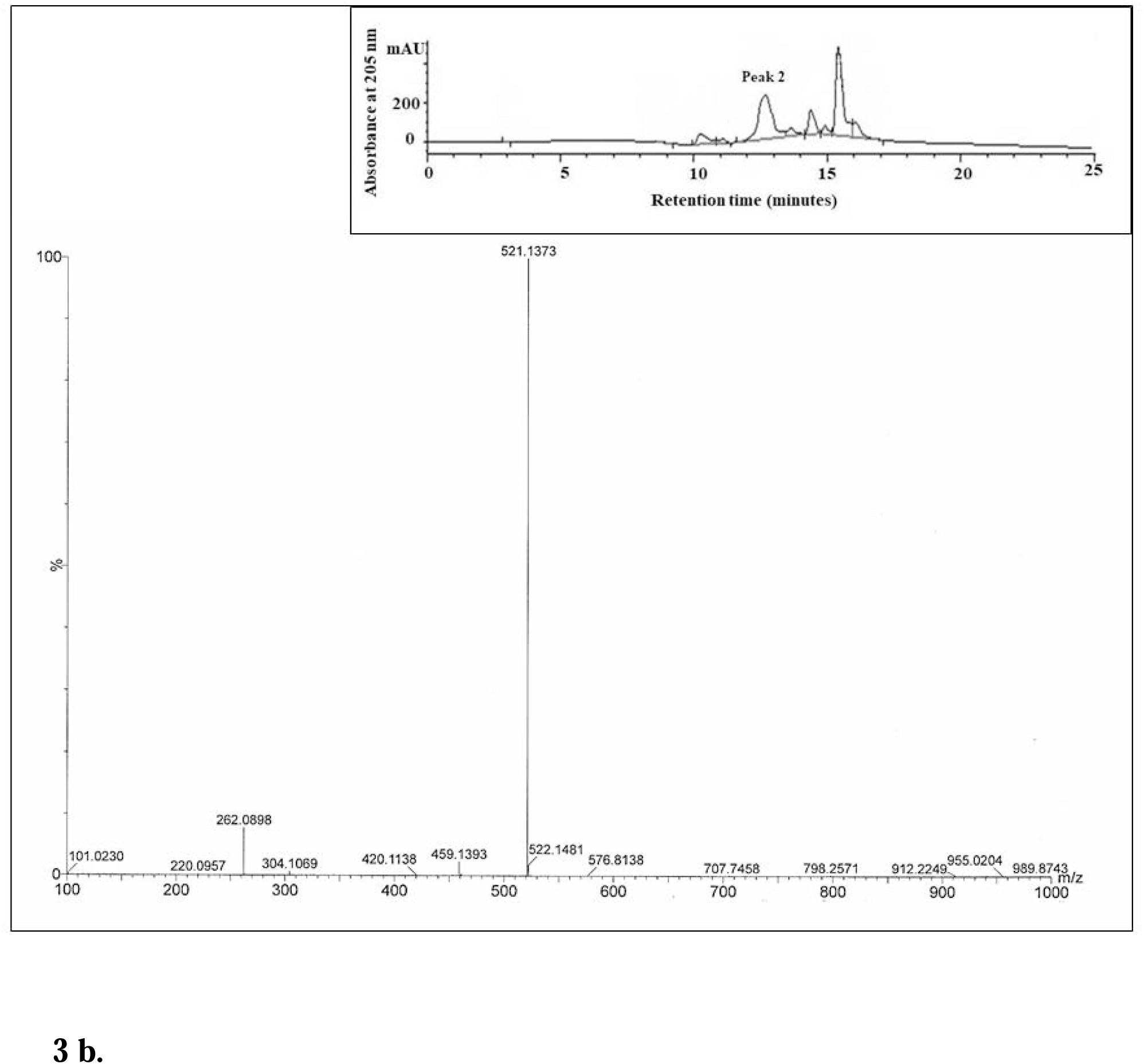
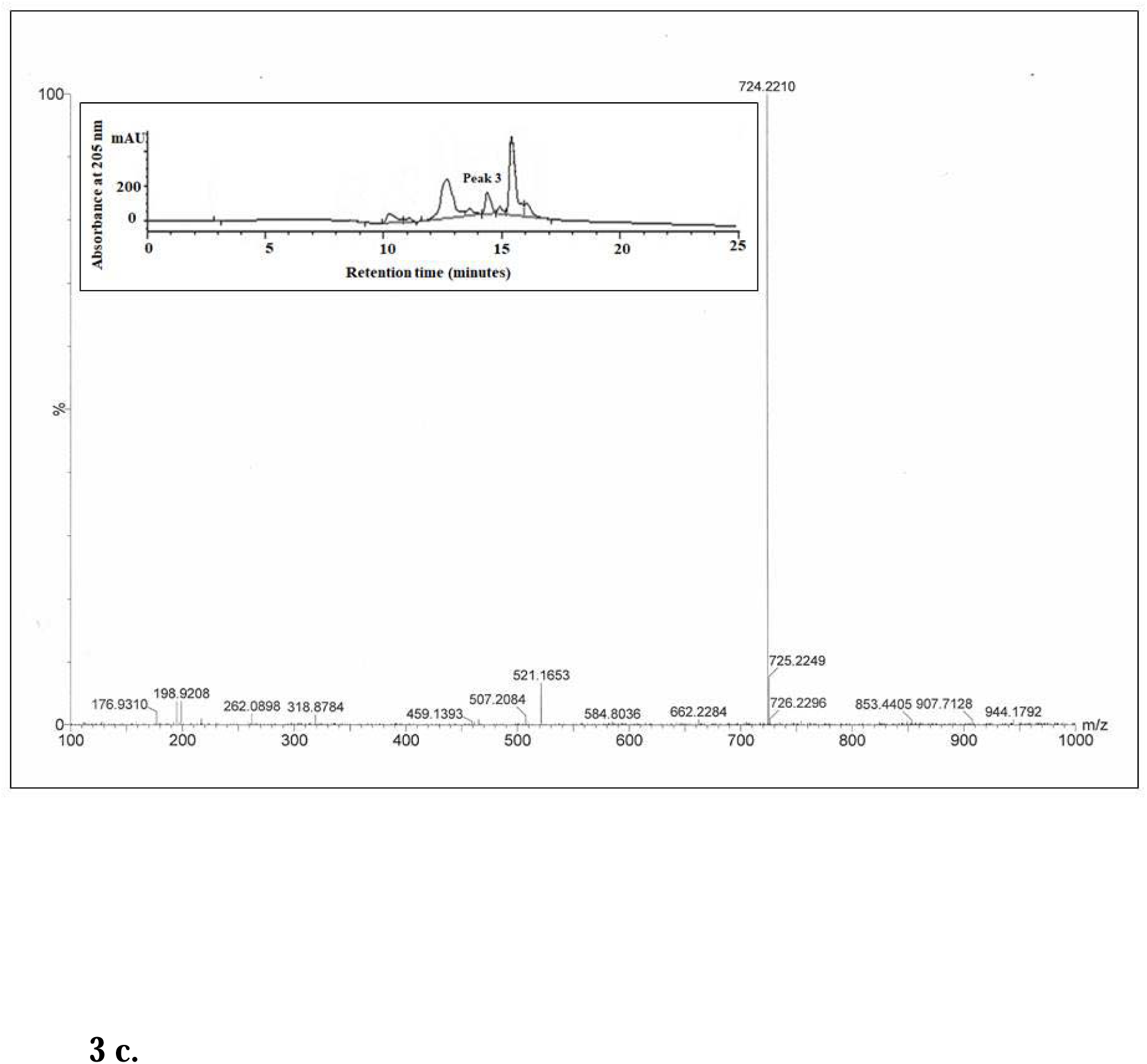
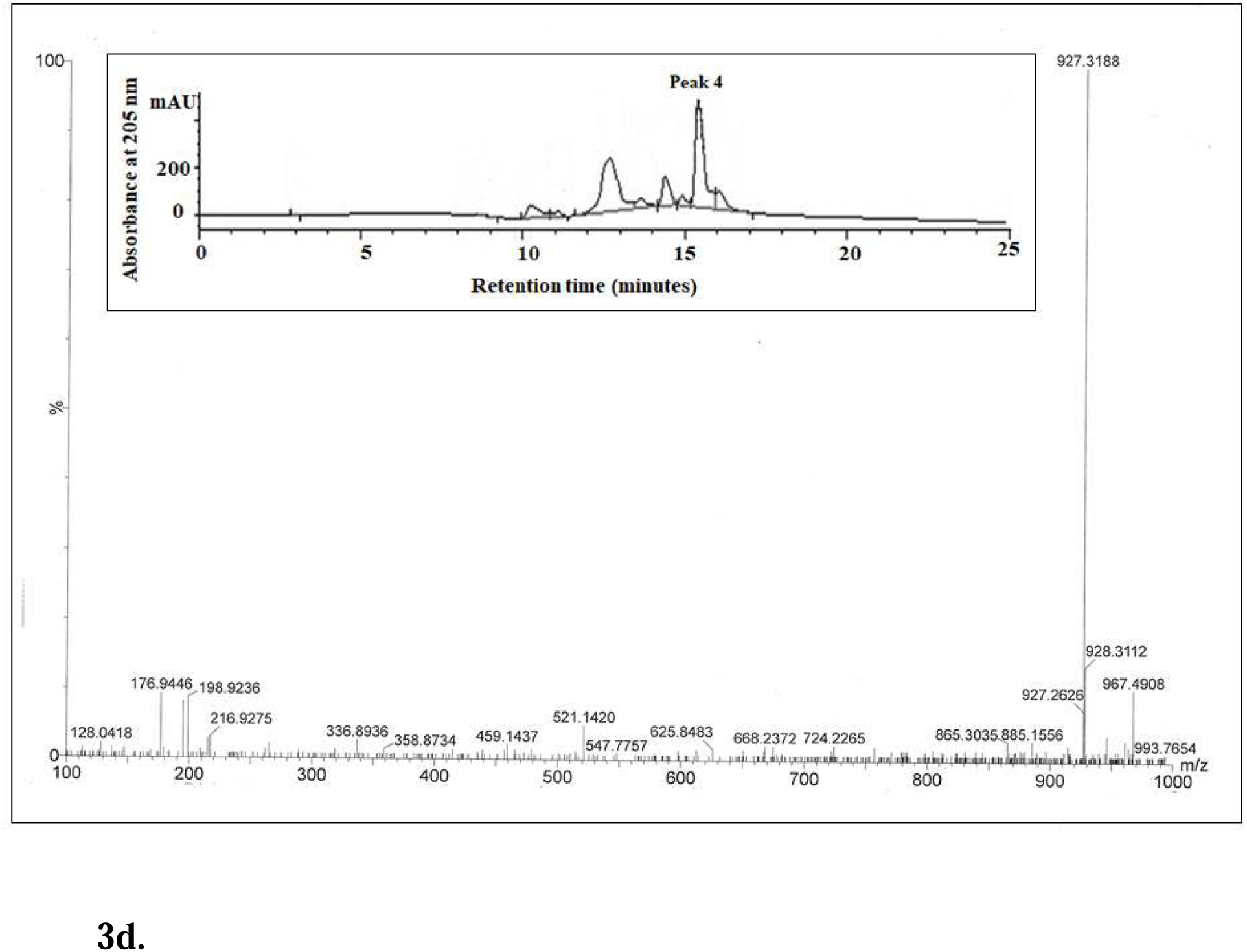
HRMS of the HPLC purified fraction of chitohexaose hydrolysed product. The data was recorded in negative ion mode of the instrument. (a) Bi sulphate (HSO ^−^) adducted GlcNAc was detected as one of the products of retention time 10.5 minutes (b) Phosphate adducted (GlcNAc)_2_ was detected as one of the products of retention time 12.5 minutes (c) Phosphate adducted (GlcNAc)_3_ was detected as one of the products of retention time 14 (d) Bi sulphate adducted (GlcNAc)_4_ was detected as one of the products of retention time 15.5 minutes (HPLC chromatogram is represented in the inset)

### 3.5. Enzyme kinetics study with different substrates

Kinetics parameters of the purified chitinase of *A. terreus* were deducted by reacting the enzyme with a chromogenic substrates *p*NPGlcNAc and a fluorogenic substrate 4-methylumbelliferyl-tri-N-acetyl-β-chitotrioside, respectively. Chromogenic substrate *p*NPGlcNAc is known to evaluate exochitinase activity of a chitinase enzyme; [20] whereas, fluorogenic 4-methylumbelliferyl-tri-N-acetyl-β-chitotrioside is widely used to study endochitinase activity [21]. Kinetic parameters of exo- and endochitinase were derived from Michaelis–Menten equation obtained by using GraphPad Prism v 5.0 software (Graph Pad Software, La Jolla, CA, USA). The purified chitinase exhibited almost 4.7fold higher catalytic efficiency with *p*NPGlcNAc as compared to 4-methylumbelliferyl-tri-N-acetyl-β-chitotrioside (Table 3). The enzyme also showed lower *Km* against *p*NPGlcNAc indicating higher affinity towards exochitinase specific substrate (Table 3).

**Table 3.**
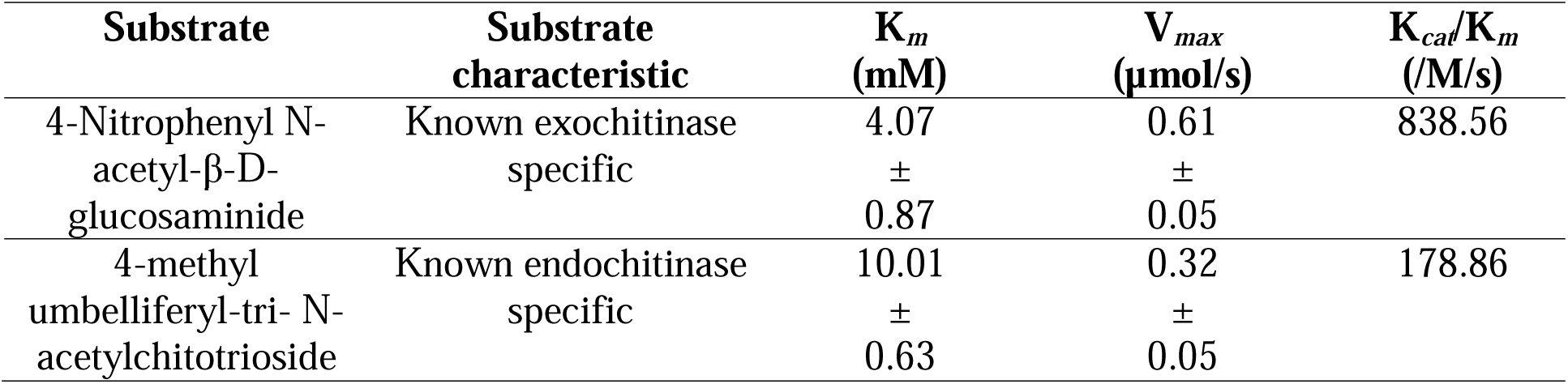
Mode of action of *A. terreus* chitinase on different artificial substrates.

| **Substrate** | **Substrate**<br>**characteristic** | **$K_m$**<br>**(mM)** | **$V_{max}$**<br>**(μmol/s)** | **$K_{cat}/K_m$**<br>**(/M/s)** |
| --- | --- | --- | --- | --- |
| 4-Nitrophenyl N-acetyl-β-D-glucosaminide | Known exochitinase<br>specific | 4.07<br>±<br>0.87 | 0.61<br>±<br>0.05 | 838.56 |
| 4-methyl<br>umbelliferyl-tri- N-<br>acetylchitotrioside | Known endochitinase<br>specific | 10.01<br>±<br>0.63 | 0.32<br>±<br>0.05 | 178.86 |

### 3.6. Subcellular localizations study of the chitinase

Amino acid sequence of the identified chitinase was extracted from NCBI database and subjected to study for subcellular localization for the protein of interest. The SignalP v4.1 determined the presence of N-terminal signal peptide containing 22 amino acids in it (Figure S2), which also confirmed the protein is translocated after translation [23]. In addition, SCL-Pred server[24] depicted the endochitinase 1 precursor of *A. terreus* to be a secretory one with high confidence level. In fact, the chitinase was originally purified from extracellular media, which is a very good agreement between *in silico* analysis and experimental observation.

### 3.7. Secondary structure analyses

Both bioinformatics tool-based prediction and circular dichroism (CD) spectroscopic analyses were carried out to demonstrate the gross conformation of the chitinase in a liquid state. The *in-silico* prediction by PSIPRED and Predict Protein demonstrated that the chitinase of *A. terreus* is comprised of 25.86% of α-Helix, 19.46% of β-sheet and 54.68% of random coil (Table 4). This was further validated by circular dichroism (CD) analyses. The characteristic CD spectra confirmed the purified protein remained stable and properly folded during purification. CD spectroscopic data of the purified chitinase was analyzed through K2D3 software, which revealed the presence of αβ characteristic structure of the protein with 28.37% of α-Helix, 19.19% of β-sheet and 52.44% of the random coil (Table 4) [25].

**Table 4.** Percentage distribution of secondary structures obtained from CD spectra and bioinformatics analysis.

| Methods of analysis | % of α-Helix | % of β-sheet | % of Random coil |
| --- | --- | --- | --- |
| Circular dichroism Analyses (K2D3 server) | 28.37 | 19.19 | 52.44 |
| Predict protein and Psipred | 25.86 | 19.46 | 54.68 |

### 3.8. Three-dimensional structure analyses

Elucidation of three-dimensional structures of marine *A. terreus* chitinase has not been carried out to date. Here, *in silico* structure analysis was chosen as the easiest alternative to laborious and time-consuming NMR or crystallography-based three-dimensional structure determinations to establish a structure-function relationship and subsequent product formation.

In the present study, *A. fumigatus* YJ-407 Chitinase (pdb id: 1WNO) was used as a template for homology model building of *A. terreus* chitinase by using Swiss Model Automated Server (Arnold et al., 2006). Here, a lower *E*-value (Expect value) was set as a parameter for homologous model searching through BLAST[33]. The final model was chosen as per QMEAN (Qualitative Model Energy Analysis) score in the Swiss Model server. The Root Mean Square Deviation (RMSD) between template and the model was observed as 0.23 Å; which indicated the present model is well derived one. Besides, pairwise alignment showed 87.9% identity between *A. terreus* and *A. fumigatus* chitinase (Figure S3). Stereochemistry and stability of the present model were assessed with PROCHECK and Verify3D respectively. According to PROCHECK analysis, 94.5% amino acid residues of *A. terreus* chitinase model were observed to present at the most favored region in the Ramachandran plot; whereas, all the residues of the model had an average 3D-1D score above 0.2, as obtained through Verify3D profile.

Like *A. fumigatus* chitinase, the three-dimensional model of *A. terreus* chitinase depicted the presence of one catalytic domain and an auxiliary α+β fold [13] (Figure 4). The catalytic domain (residues 44-297; 364-404) was found to be composed of eight-stranded α /β barrel (TIM barrel) and located at the N-terminal region of the chitinase. In this catalytic domain, different aromatic and acidic residues (Asp and Glu) were observed to be present which are presumed to form the sub-sites for the substrate binding and catalysis [34,35]. The 297-363 was found to be adopted one α+β fold, which seemed to be formed by an insertion in the barrel motif (Figure 4)[36].

**Figure 4.**
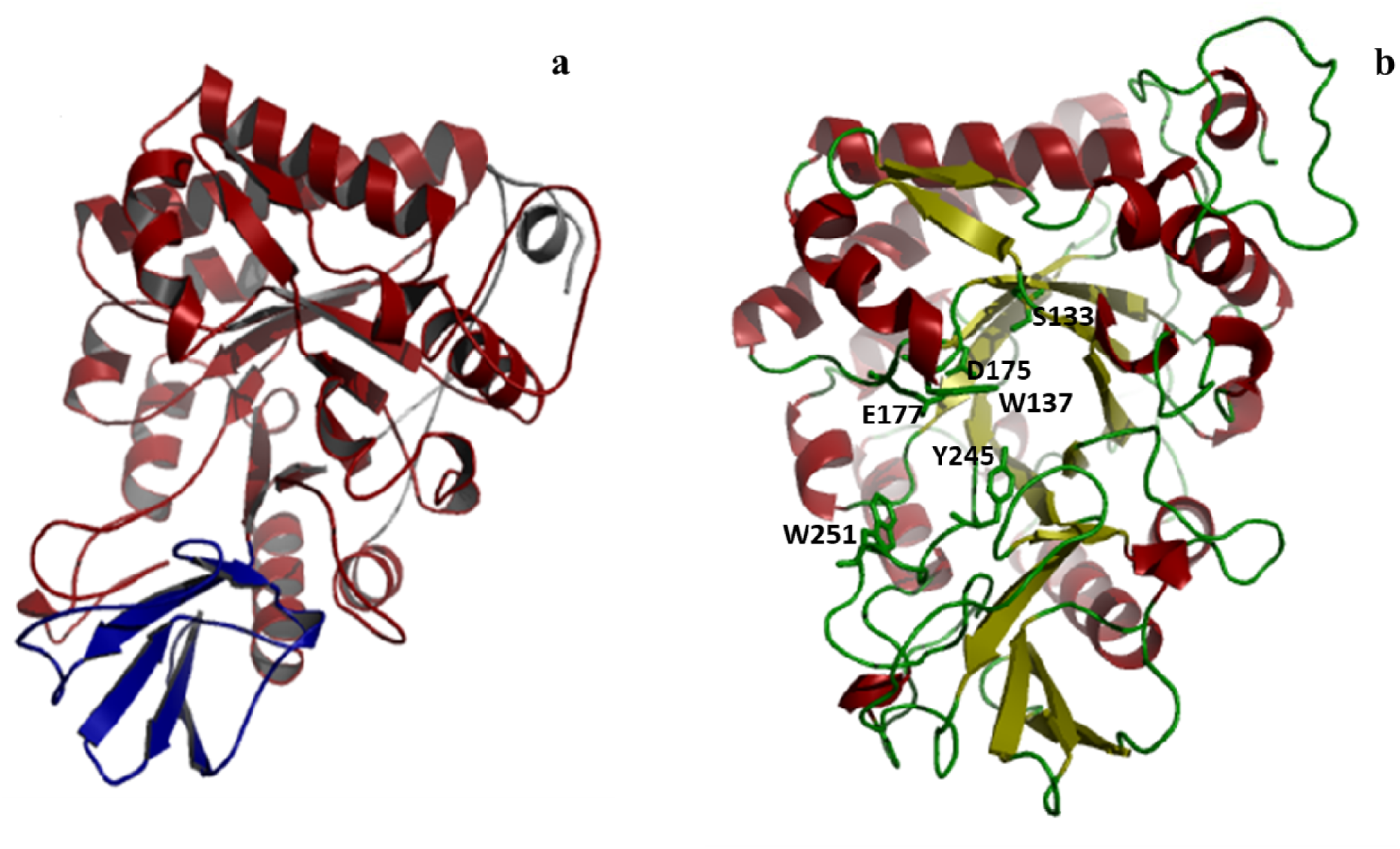
(a) SWISS Prot derived Ribbon diagram of *A. terreus* chitinase. Here, two domains are coloured differently; the α/β-barrel domain in red and the α+β domain in blue.(b) Important residues are labelled in the chitinase model

Multiple sequence analyses of the *A terreus* chitinase also revealed the presence of two conserved consensus motif boxes (Box 1 and Box 2) at the N-terminal region of the protein (Figure 5), which was characteristic of family 18 chitinases [4]. The chitin hydrolysis domain in conserved regions contained serine and glycine residues in Box 1, and glutamic acid (E) and aspartate (D) residues in Box 2 (Figure 6). From the earlier study, it can be predicted that Asp 175 and Glu 177 of DxxDxDxE motif at the end of β4 might possibly serve in the substrate binding [13] (Figure 5); while Glu 177 was presumed to serve as catalytic acid that protonates the scissile glycosidic bond [37]. In course of chitin hydrolysis, Glu 177 was proposed to contribute carboxyl hydrogen to form oxazolinium ion as intermediate[35] whereas; stabilization of the intermediate by ion pairing with the developing positive charge was predicted to be performed by Asp 175 and subsequent lowering the pKa of Glu 177 (Figure 5)[38]. Importance of these residues had been established by site-directed mutagenesis of the corresponding residues in Chit42 of *T. harzianum* (Glu172 and Asp170 of *T. harzianum*)[5]. Comparing the structural insight into catalytic mechanism of a family 18 chitinase of *Serratia marcescens*, it was predicted that Asp 173 in *A. terreus* chitinase might be involved in rising pKa of Asp175 while, Ser 133 of SxGG motif might stabilize Asp 175 during catalysis[35].

**Figure 5.**
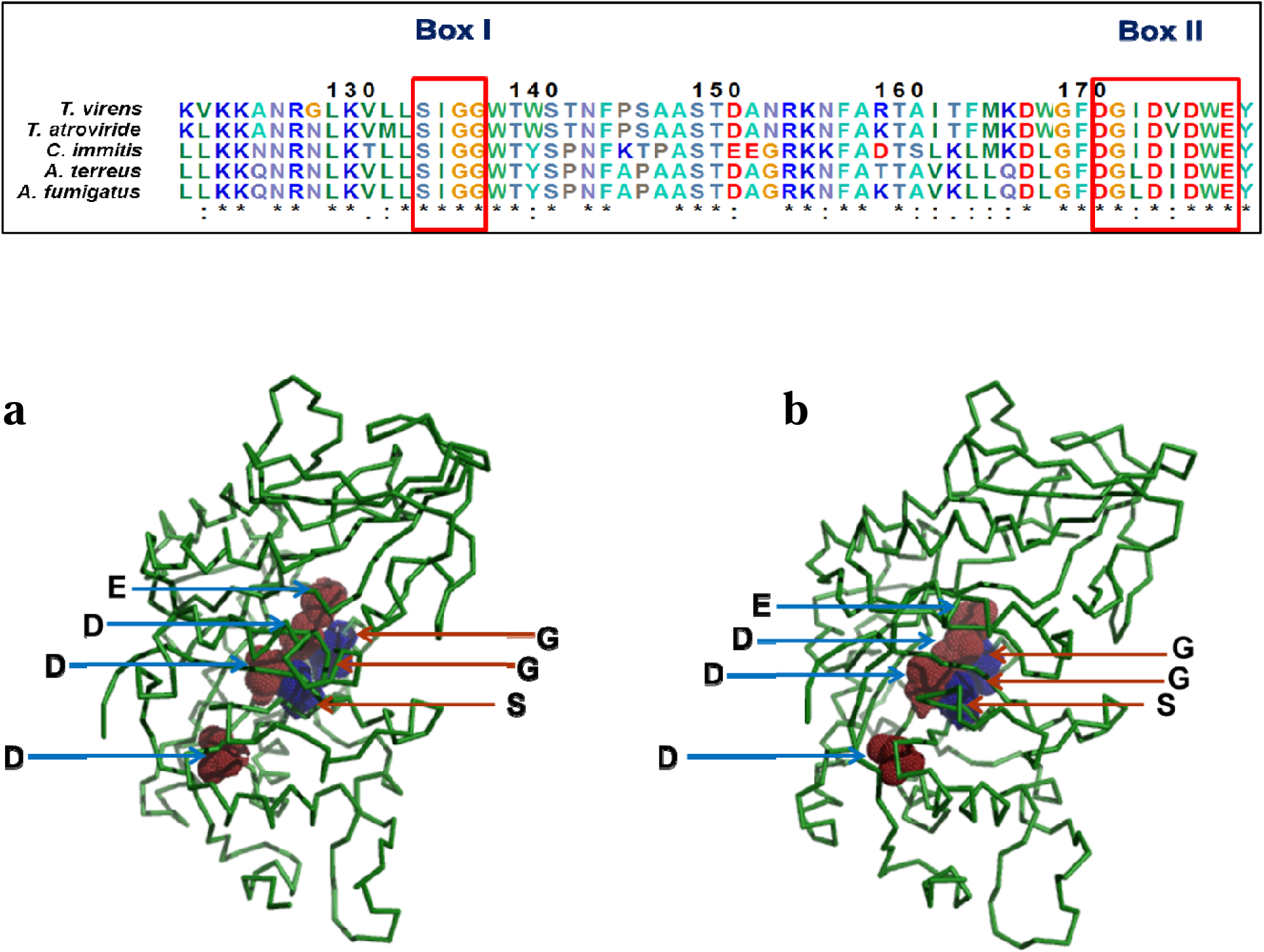
Location of SxGG (Box 1) and DxxDxDxE (Box 2) motifs in the structure of (a) chitinase from *A. fumigatus* (b) chitinase of *A. terreus* (Comparison of active site residues of different fungal chitinases by Clustal omega analysis in box)

From the earlier literature it was predicted that the conserved Trp 137 and Trp 251 of *A. terreus* chitinase had probably undergone conformational changes to create a hydrophobic sandwich with the sugars and helped in enzyme-substrate interaction like othe family 18 chitinases[35]. In present study, *A. terreus* chitinase showed a dual mode of action like *the* homologous counterpart of *A. fumigatus* [7]; which is confirmed by both molecular docking along with biochemical study of the present chitinase with exochitinase and endochitinase specific artificial substrates. The active site of *A. terreus* chitinase was observed to possess a deep tunnel shape with the number of aromatic residues for proper binding of the chitin polymer (Figure 5). The aromatic residues form subsites might have a role in interacting hydrophobically with GlcNAc moiety of the chitin chain. In the present study, the best-fitted ligand receptor complex was selected on the presence of active site-specific amino acids in CB Dock 2 based blind docking derived complexes. The molecular docking of the present chitinase showed the lower binding energy for exochitinase-specific substrate in comparison to endochitinase-specific one (Table 5); which further confirm not only the dual mode of action of this chitinase but also higher catalytic activity toward exo-chitinase specific behavior as derived by enzyme kinetics analyses by biochemical assay.

**Table 5:** CB Dock 2 derived docking score of ligands with *A. terreus* chitinase.

| Receptor | Template Protein id | Ligand | Fit Doc Score | Interacting amino acids |
| --- | --- | --- | --- | --- |
| Chitinase of *Aspergillus terreus* | 4wkf | *p*-Nitrophenyl N-acetyl-β-D-glucosaminide | -3.5 | TRP137 GLU177<br>TYR178 ASP246<br>ARG301 GLU322<br>TRP384 |
| | | 4-methyl umbelliferyl-tri-N-acetyl-β-chitotrioside | 1703.2 | TRP137 GLU177<br>TYR178 LYS224<br>TYR245 ASP246<br>TRP251 ARG301<br>GLU322 TRP384 |

## 4. Discussion

The production of GlcNAc and other derivatives from chitin polymer relies on the enzymatic property and mode of action of chitinase [1,2,9]. Several researchers have explored different microbial chitinases to valorise the chitin wastes and subsequent production of GlcNAc or its multimeric derivatives for their potential health care and environment-related applications [[1,2,9]. However, very few potential microbial chitinases have been systematically characterized to investigate their chitin hydrolysis mechanism[9,39]. Here, chitin hydrolysis pattern of a marine *A. terreus* chitinase has been illustrated with the help of different biochemical assays, biophysical characterizations and bioinformatics analyses to unveil the key mechanism for GlcNAc production from chitin as well as classify the present chitinase as per conventional classification pattern.

Earlier literatures described that the hydrolysis pattern is the key basis of classification of chitinases, which can be determined by the nature of product released [7]. An exochitinase excises the chitin polymer from non-reducing terminal end and results either GlcNAc or its dimer (diacetylchitobiose)[7,40] . On the other hand, an endochitinase cleaves chitin chain randomly and produce N-acetyl-chitooligosaccharide derivatives of different length [7,40]. In this study, HPLC followed by High Resolution Mass Spectrometric (HRMS) analyses were performed to characterize the product and the nature of the isolated chitinase from *A. terreus*. The enzymatic action of swollen chitin polymer released only GlcNAc as a sole product as detected by HPLC and HRMS, even after varying the incubation period, indicating an exochitinase behaviour of the purified chitinase. Notably, the residual polymer was not detected with the method followed by in this study. We assume that since only the soluble fraction of the hydrolysate was injected into HPLC column after centrifugation, the insoluble remainders might have been filtered out. By contrast, with hexa-N-acetylchitohexaose, both GlcNAc and N-acetyl-chitooligosaccharides of varied lengths were detected from hydrolysates. Whilst the isolated chitinase exhibited predominantly exochitinase activity against natural substrate, the endochitinase activity has become evident with the substrate with smaller chain length. This observation of dual characteristic of purified enzyme is however in contrary to its molecular identification, as revealed from peptide mass finger printing analyses by MALDI Tof/Tof, which identified purified chitinase as endochitinase 1 precursor. Notwithstanding the annotation, no other chitinase with similar mass was identified suggesting only single enzyme was purified and tested in this study.

To further investigate the mechanism of enzymatic action, the kinetic studies with a dimeric chromogenic and a tetrameric fluorogenic substrates were carried out with the purified enzyme of *A. terreus*. With 4-Nitrophenyl N-acetyl-β-D-glucosaminide or *p*NPGlcNAc, a monomeric GlcNAc and chromophoric *p*-nitrophenol were produced, clearly indicating exochitinase behaviour of the enzyme. Whereas, with 4-methylumbelliferyl-tri-N-acetyl-β-chitotrioside, a trimer of GlcNAc and a fluorescent compound 4-methylumbelliferone were produced, indicating endochitinase activity of the enzyme. Surprisingly, catalytic efficiency of enzyme with dimeric substrate is higher than tetrameric substrate. Precedence for dual nature has been evidenced with a number of family 18 chitinases for a few fungal species like *A. fumigatus* and *Penicillium oxalicum* k10 [7,15], archaea like *Thermococcus kodakaraensis* KOD1 [40,41] and a marine bacterium *Saccharophagus degradans* (formerly *Microbulbifer degradans*) [42]. However, the current study at the best of our knowledge is the pioneer to report the dual nature of a family 18 chitinase from a marine *Aspergillus terreus*.

Other than fungal chitinases, the family 18 chitinases of some archaea and bacteria were reported to have dual GH18 catalytic domains [40,41]. Mutagenesis study and individual domain analyses revealed differential catalytic behaviours of the two domains accounting for endo- and exochitinase activity of these organisms[40,41].

On the contrary, the isolated chitinase from *A. terreus* and its homologous counterpart from *A. fumigatus* belong to GH18 family and possess single catalytic domain. Differential cleavage specificities of the single domain chitinases upon varied length of the substrate is enigmatic; which was compared with an earlier study on a classical structural experiment with *Serratia marcescens* chitinase A complexed with the chito-oligosaccharides of varying size and length explained different cleavage pattern due to different mode of substrate binding to the aromatic sub-sites[4] . Although, the structural aspect of the dual activity of the isolated chitinase from *A. terreus* is not properly elucidated yet however, it can be speculated that accessibility of the substrate towards binding residues through the tunnel-shaped binding cleft possibly depends on the substrate length, which renders different processive hydrolysis mechanism. Further, lower binding energy requirement of *A. terreus* chitinase for exochitinase-specific substrate establishes its predominant characteristic as exochinase in contrust to endochitinase one. Therefore, it can be more likely to presume why the isolated chitinase of *A. terreus* exhibited higher affinity and catalytic efficiency towards smaller dimeric substate as compared to tetrameric substrate.

Intriguingly the isolated chitinase from *A. terreus* exhibited its stability at higher temperature, which may open up broad applicability of the enzyme in several biocatalytic industries. Nevertheless, the structural basis of stability of the isolated chitinase from marine *A. terreus* awaits experimental substantiation.

The current study would render a basis of understanding the functional characterization of chitinase from marine *Aspergillus terreus* fungal strain and provide deeper insight into the link between chitin degradation and substrate length; which may reveal new mechanistic insight on future physiological and biochemical investigations.

### Conclusions

The purified chitinase of marine *A. terreus* identified as endochitinase 1 precursor was perhaps mere annotation in NCBI database, without any experimental evidence. In the present study, biochemical basis of the nomenclature and relationship between three-dimensional structure and mechanism of action were investigated. Mechanism of different substrate conversion by *A. terreus* chitinase was established in the present study. Here, MALDI ToF/ToF study identified the purified enzyme as an endochitinase precursor, whereas, GlcNAc and its oligomer production attributes the dual nature of the isolated enzyme, as determined by HRMS analyses. The isolated chitinase has demonstrated predominant exochitinase activity over its endochitinase action, while studying its kinetics in the presence of specific artificial substrates. The *in silico* structural analyses of the chitinase also corroborated the structural basis of its dual mode of action and hence, strongly established the hypothesis that this chitinase of marine fungal origin could qualify as an efficient biocatalyst for potential commercial production of GlcNAc from most of the known natural sources of chitin.

## Supplementary information

E-supplementary data of this word can be found in online version of the paper.

## CRediT authorship contribution statement

**Sancharini Das:** Writing original draft, Data curation, Formal analysis, Investigation, Methodology, Writing - review & editing; Validation, Resources **Debasis Roy:** Investigation; Methodology; Project administration, Supervision; Resources Writing - review & editing **Ramkrishna Sen**: Conceptualization; Investigation; Methodology; Project administration, Supervision; Resources Writing - review & editing

## Acknowledgment

SD is thankful to National Jute Board (NJB), Ministry of textile Govt. of India, IIT Kharagpur for Research Assistantship and DBT BioCARe Programme (Project Code: BT/PR50872/BIC/101/1294/2023) for financial and academic support for the research work. Dr. Ramapati Samanta and Dr. Chiranjit Chowdhury is gratefully acknowledged for HPLC related and bioinformatics related research work respectively.

